# Inhibition of Androgen Receptor Exposes Replication Stress Vulnerability in Prostate Cancer

**DOI:** 10.1101/2024.10.08.617102

**Authors:** Carly S. Williams, Xin Li, Hongjun Jang, Jay Ramanlal Anand, Won Young Lim, Hyejin Lee, Julie Parks, Xingyuan Zhang, Jialiu Xie, Jinshi Zhao, Di Wu, Andrew J. Armstrong, Jessica L. Bowser, Lee Zou, Jiyong Hong, Jason A. Somarelli, Cyrus Vaziri, Pei Zhou

## Abstract

Standard initial systemic treatment for patients with metastatic prostate cancer includes agents that target androgen receptor (AR) signaling. Despite an initial positive response to these AR pathway inhibitors (ARPIs), acquired resistance remains a significant challenge. We show that treatment of AR-positive prostate cancer cells with the frontline ARPI enzalutamide induces DNA replication stress. Such stress is exacerbated by suppression of translesion DNA synthesis (TLS), leading to aberrant accumulation of single-stranded DNA (ssDNA) gaps and persistent DNA damage biomarkers. We further demonstrate that the TLS inhibitor, JH-RE-06, markedly sensitizes AR-positive prostate cancer cells, but not AR-negative benign cells, to enzalutamide *in vitro.* Combination therapy with enzalutamide and JH-RE-06 significantly suppresses cancer growth in a syngeneic murine tumor model over vehicle control or individual treatment groups. These findings suggest that AR inhibition broadly triggers DNA replication stress in hormone-sensitive prostate cancer, thereby exposing a unique vulnerability that can be exploited by a TLS-disrupting adjuvant for targeted therapy.

## INTRODUCTION

Prostate cancer is the second leading cause of cancer mortality in men (*1*). It is estimated that 1 in 8 men will be diagnosed with prostate cancer within their lifetime, and about one in 44 men will die of prostate cancer in the US (*2, 3*). Androgen receptor pathway inhibitors (ARPIs), such as enzalutamide, apalutamide, abiraterone acetate and darolutamide, are effective first-line therapies for men with metastatic prostate cancer, prolonging patient survival (*4–10*). However, resistance to ARPIs in men with metastatic hormone-sensitive prostate cancer develops typically within several years despite an initial favorable response, and thus overcoming or delaying acquired ARPI resistance and castration resistance is a major unmet medical need (*4–10*).

Blockage of AR significantly alters the transcriptional profile of numerous genes, including homologous recombination repair (HRR) and other DNA repair genes (*11–14*). Compromised DNA repair activities, including defective HRR, frequently leads to slowing or stalling of DNA replication forks, a phenomenon known as “replication stress” that renders cells heavily reliant on translesion DNA synthesis (TLS) to resolve the stress and complete genomic replication (*15–17*). However, the consequence of AR blockage during genomic replication has not been investigated. Using a combination of DNA fiber assays, biomarker analysis, fluorescence-activated cell sorting (FACS), and live cell imaging, here we provide the first experimental evidence that enzalutamide treatment induces replication stress in prostate cancer, which is efficiently resolved by TLS, resulting in reduced replication fork progression without accumulation of ssDNA gaps. Simultaneous treatment of prostate cancer cells with enzalutamide and the TLS inhibitor JH-RE-06 (*18*) not only slows DNA replication, but also leads to accumulation of single stranded DNA (ssDNA) gaps, increased levels of DNA replication stress biomarkers, and reduced mitotic entry. Accordingly, JH-RE-06 profoundly sensitizes AR-positive prostate cancer cells to enzalutamide treatment both *in vitro* and in a syngeneic murine allograft tumor model. These findings suggest that blockage of AR broadly induces replication stress in hormone-sensitive prostate cancer, exposing a unique vulnerability that can be exploited by TLS-targeting adjuvant cancer therapeutics.

## RESULTS

### Enzalutamide treatment induces DNA replication stress

Enzalutamide treatment of prostate cancer cells was reported to suppress homologous recombination repair (HRR) genes (*11, 12*). Consistent with these reports, our own RNA-Seq analysis revealed a significant perturbation of the transcription profile of DNA damage response (DDR) genes in enzalutamide-treated TRAMP-C2 cells in comparison with the DMSO control group (**Fig. S1**). However, the molecular consequences of enzalutamide treatment in the context of genomic replication has not been established. To address this knowledge gap, we used 5-Bromo-2’-deoxyuridine (BrdU) labeling and FACS to measure the rates of DNA replication in 22Rv1 and TRAMP-C2 prostate cancer cells treated with enzalutamide or DMSO (negative control) (**Fig. 1A**). Although treatment of TRAMP-C2 cells with 25 μM enzalutamide showed no effect in comparison with the DMSO control, treatment of 22Rv1 cells with 50 μM enzalutamide led to a ∼24% decrease in rates of BrdU incorporation, potentially indicating that the AR antagonist enzalutamide induced DNA replication stress. Since TLS often plays a critical role in mitigating the DNA replication stress (*16, 17*), we investigated the consequence of TLS inhibition by JH-RE-06 (*18*) on enzalutamide-induced DNA replication stress. As shown in **Fig. 1A**, treatment of 22Rv1 cells with 5 μM JH-RE-06 did not reduce rates of DNA synthesis in the absence of enzalutamide; however, combined treatment with 50 μM enzalutamide and 5 μM JH-RE-06 led to a 73% decrease in BrdU incorporation rates of 22Rv1 cells relative to the DMSO control cultures (**Fig. 1A**). A similar synergistic effect to reduce the DNA synthesis rates was also observed in TRAMP-C2 prostate cancer cells treated with 25 μM enzalutamide and 2 μM JH-RE-06 (**Fig. 1A**). In both 22Rv1 and TRAMP-C2 cell lines, the combined treatment with enzalutamide and JH-RE-06 additionally induced a population of BrdU-positive cells with late S-phase /G2 DNA content (indicated by pink dashed line on far-right panels of **Fig. 1A**). This result suggests that DNA synthesis persists aberrantly late in the cell cycle when TLS fails to resolve enzalutamide-induced replication stress arising from early and mid S-phases.

**Figure 1.**
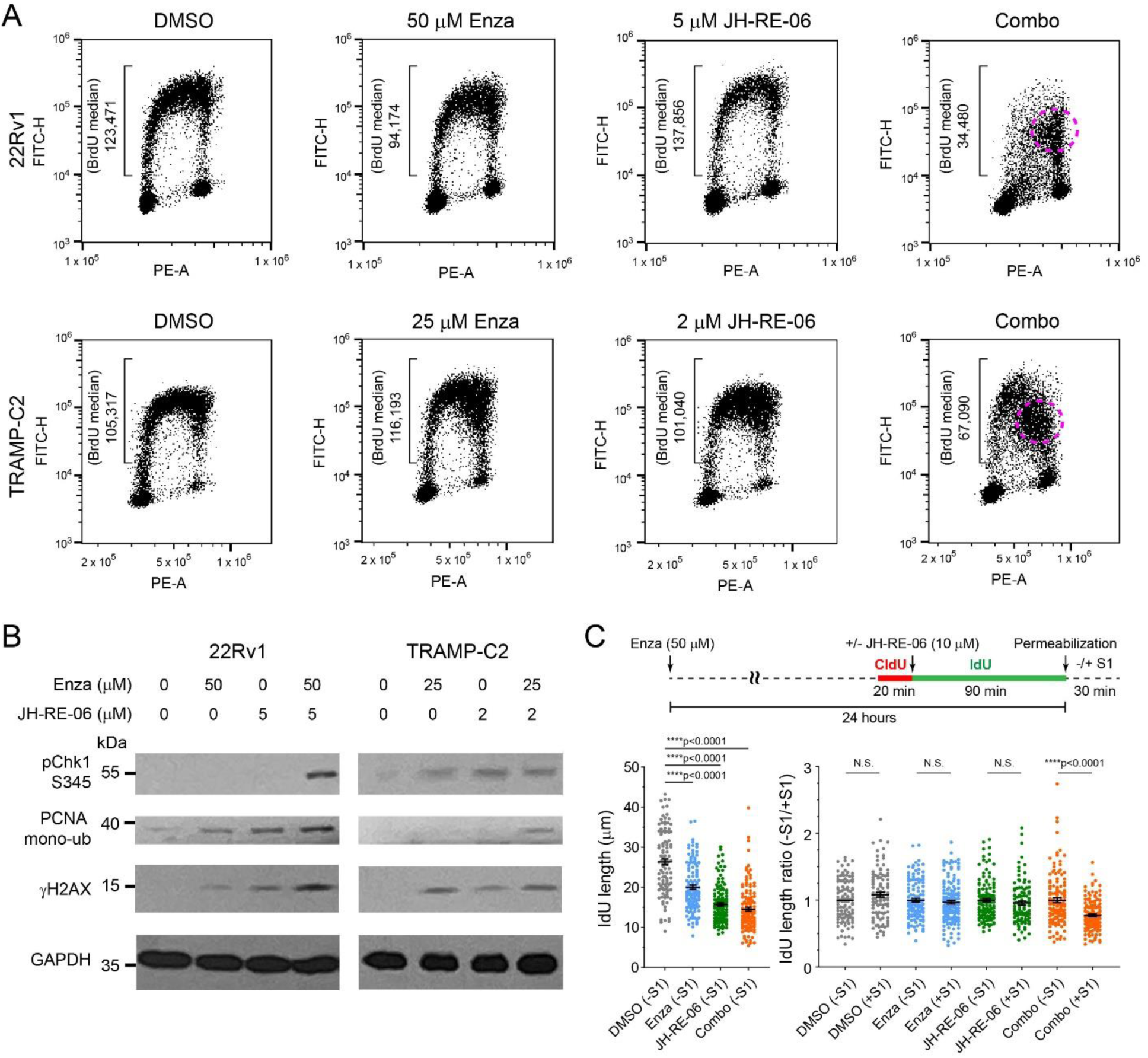
Treatment of prostate cancer cells with enzalutamide induces replication stress that is amplified by inhibition of translesion DNA synthesis. (A) Representative FACS data of 22Rv1 and TRAMP-C2 cells treated with DMSO, enzalutamide (Enza), JH-RE-06 and their combination. In the combination treatments, 50 μM enzalutamide and 5 μM JH-RE-06 were used for 22Rv1 and 25 μM enzalutamide and 2 μM JH-RE-06 were used for TRAMP-C2 cells. Replication stalling during the S-to-G2 transition is indicated by dashed circles colored in pink. (B) Western blotting of DNA repair biomarkers of 22Rv1 and TRAMP-C2 cells treated with the AR inhibitor enzalutamide and the TLS inhibition, JH-RE-06. (C) DNA fiber assays of 22Rv1 cells treated with DMSO, 50 μM enzalutamide (Enza), 10 μM JH-RE-06, and their combination. The lengths of IdU labeled DNA fibers of 22Rv1 cells in different treatment groups without the S1 nuclease treatment are shown in the left panel, and the length ratio of S1 nuclease treated DNA fibers over their untreated counterparts are shown in the right panel. Each dot represents one fiber. At least 100 fibers are measured from two independent experiments. Error bars represent S.E.M. Statistical analysis: one-way ANOVA followed by Brown-Forsythe and Welch multiple comparison test. ****P < 0.0001.

Consistent with our cell cycle analyses, we found that simultaneous inhibition of AR and TLS in 22Rv1 or TRAMP-C2 cells significantly and consistently elevated the levels of phospho-Chk1 (pChk1, a marker of the replication stress response), mono-ubiquitinated PCNA (required for TLS), and γH2AX (a general marker of DNA damage), indicating that the combination treatment significantly induced replication stress when both AR and TLS are inhibited (**Fig. 1B**). Individual treatment with either enzalutamide or JH-RE-06 also increased the levels of DDR biomarkers indicated above relative to the DMSO control group; however these changes were not uniform in the two tested cell lines (22Rv1 and TRAMP-C2), and they were generally more modest in comparison with the combination treatment (**Fig. 1B**). Although heterogeneous responses to targeted therapies (as observed in our single agent case) are frequently encountered in distinct cancer cell lines due their varied repertoires of resistance mechanisms, such as the presence of AR splice variants in 22Rv1 cells (*19*), it is encouraging that both of these cells lines responded uniformly with biomarkers to the combination treatment, indicating a broadly-effective outcome.

Since reduced levels of BrdU incorporation in our FACS assay can be potentially caused by reduced origin firing or from slowing/stalling of fork movement as in replication stress, we measured the rates of DNA synthesis at the resolution of individual forks using DNA fiber assays to formally test whether enzalutamide and JH-RE-06 treatments cause replication stress (**Fig. 1C** & **Fig. S2**). Briefly, 22Rv1 prostate cancer cells were treated with the AR inhibitor enzalutamide (50 μM) for ∼24 hours, and the nascent DNA was labeled with the thymidine analog 5-chloro-2′-deoxyuridine (CldU) for 20 minutes followed with 5-iodo-2′-deoxyuridine (IdU) for 90 minutes. The consequences of TLS inhibition by JH-RE-06 and its combination with enzalutamide were evaluated by adding JH-RE-06 (5 μM) simultaneously at the beginning of the IdU labeling period. Post labeling, cells were permeabilized and treated either without or with the S1 nuclease to detect the presence of ssDNA gaps, as cleavage of ssDNA would reduce the length of DNA fibers containing these gaps. We found that IdU-labeled DNA fibers in enzalutamide-treated 22Rv1 cells were significantly shorter than those in the DMSO control group, reflecting slowing or stalling of the replication fork progression; similar phenomena were observed for 22Rv1 cells treated with the TLS inhibitor, JH-RE-06, or the combination of enzalutamide and JH-RE-06 (**Fig. 1C**, left panel). Importantly, our observation of a dramatic replication fork slowdown in enzalutamide-treated cells supports the notion that enzalutamide treatment alone induces replication stress. In this regard, it is particularly interesting to note that upon the treatment of the S1 ssDNA nuclease, the only significant length reduction was observed in the enzalutamide/JH-RE-06 combination treatment group, but not in the DMSO control or individual treatment groups of enzalutamide or JH-RE-06 (**Fig. 1C**, right panel; **Fig. S2**). Considering the elevation of monoubiquitinated PCNA, an event involved in TLS, in enzalutamide-treated 22Rv1 cells over the DMSO control group (**Fig. 1B**), it appears that ssDNA gaps resulting from enzalutamide-induced replication stress were efficiently filled by the elevated TLS activity, whereas suppression of TLS by JH-RE-06 prevented gap repair and exposed these ssDNA gaps generated from the enzalutamide-induced replication stress.

### Live cell imaging reveals defective cell cycle under the enzalutamide/JH-RE-06 combination treatment

Persistent DNA damage caused by unresolved DNA replication stress often leads to defects in G2/M progression, prompting us to examine how the combination treatment affected prostate cancer cell growth using live cell imaging. To do this, cells were transduced with lentiviruses encoding a GFP-tagged histone 2B to track mitotic events in real time. Asynchronous GFP-H2B-expressing TRAMP-C2 cells were incubated with DMSO (negative control), enzalutamide (10 μM), JH-RE-06 (2 μM), and their combination for 12 hours before being placed under the microscope for continuous image collection every three minutes for a total of 24 hrs. We found that TRAMP-C2 cells treated with either 10 μM enzalutamide or 2 μM JH-RE-06 showed a modest increase in the duration of the interphase (**Fig. 2A,B**), a phenotype consistent with the observation of slowed DNA replication in the DNA fiber assay of 22Rv1 cells treated with either enzalutamide or JH-RE-06 (**Fig. 1A**). Treatment with 10 μM enzalutamide additionally caused a small increase in the proportion of cells that displayed pre-mitotic defects in the interphase and cells that never divided, though neither reached statistical significance (**Fig. 2C,D**).

**Figure 2.**
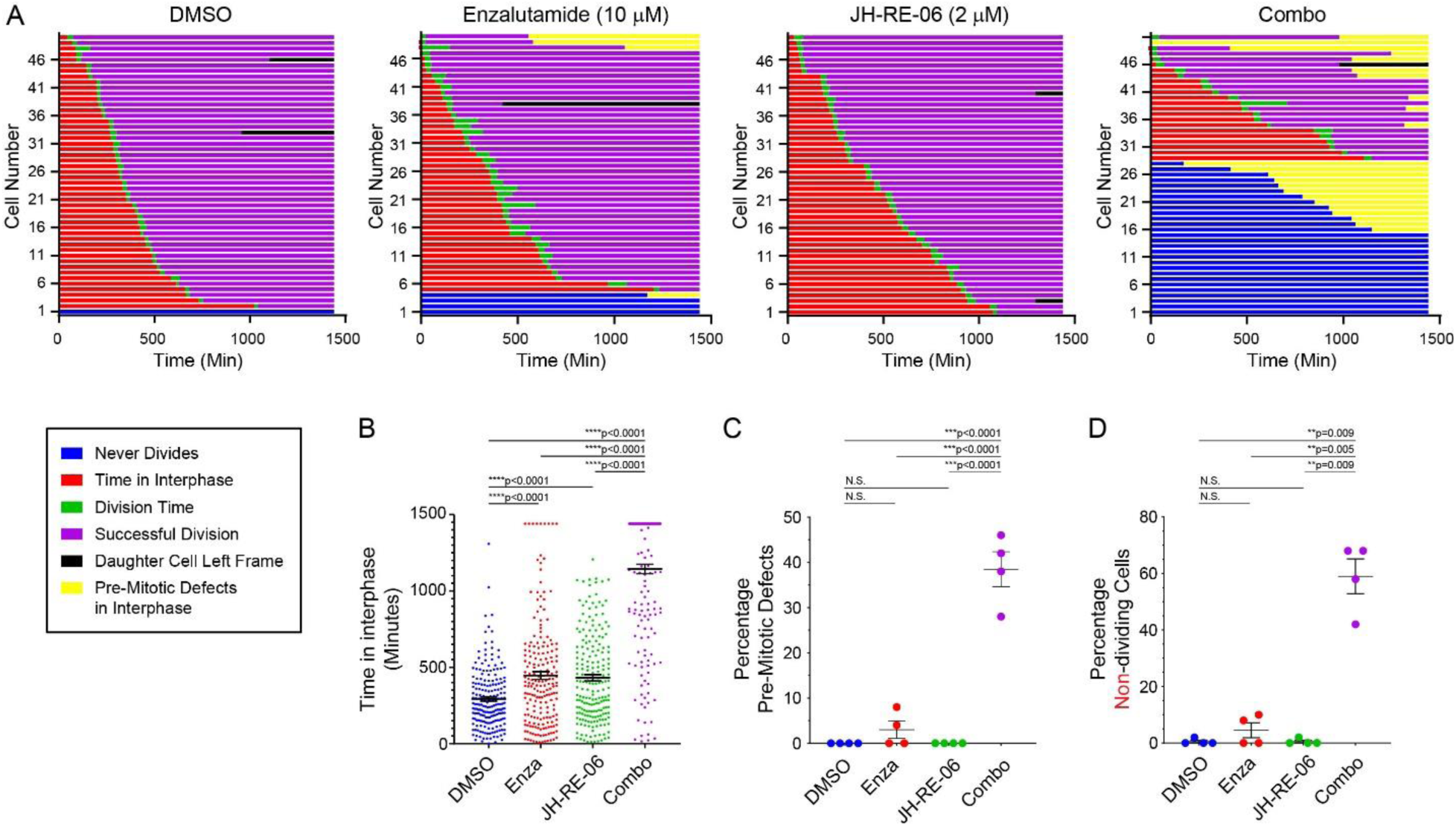
Live-cell imaging of cell cycle progression upon combined exposure to JH-RE-06 and enzalutamide. (A) Representative cell fate analysis of TRAMP-C2 after 24-hour treatment with either DMSO, enzalutamide (Enza, 10 μM), JH-RE-06 (2 μM), or their combination. Experiments were performed in technical quadruplicates, counting 50 cells per repeat. Time of individual cells in interphase, percentage of cells experiencing pre-mitotic defects, and percentage of non-dividing cells under different treatment conditions are shown in panels (B), (C), and (D), respectively. Error bars represent S.E.M. Statistical analysis: (**B**, **D**) one-way ANOVA followed by Brown-Forsythe and Welch multiple comparison test; (**C**) one-way ANOVA; ****p<0.0001; **p<0.01.

Strikingly, all of these modest phenotypical changes in enzalutamide-treated cells were dramatically amplified in enzalutamide/JH-RE-06 co-treated cells. Cells treated with a combination of enzalutamide and JH-RE-06 showed a significantly prolonged interphase with the mean duration time almost quadrupling relative to the DMSO control group and tripling relative to the individual treatment groups of enzalutamide and JH-RE-06 (**Fig. 2B**; p<0.0001 for combination treatment versus all other groups). The extended interphase of enzalutamide and JH-RE-06-treated cells is consistent with the reduced rates of BrdU incorporation and the reduced fork velocities we observed under these treatment conditions (**Fig. 1A**). About 40% of the enzalutamide/JH-RE-06 co-treated cells showed significant pre-mitotic defects in the interphase (**Fig. 2C**; p<0.0001 for combination treatment versus all other groups), and more than half of the cells never divided (**Fig. 2D**; p<0.01 for combination treatment versus all other groups). Taken together, our live cell imaging experiments revealed a pronounced mitotic defect in the enzalutamide/JH-RE-06 co-treated cells, likely due to the persistence of S-phase-dependent DNA damage induced by the combination treatment.

### Inhibition of TLS selectively sensitizes AR-positive prostate cancer cells to enzalutamide treatment

Given our observation of enzalutamide-induced replication stress, involvement of TLS in stress resolution, and defective cell cycle under the enzalutamide/JH-RE-06 combination treatment, we investigated whether TLS inhibition by JH-RE-06 could broadly sensitize AR-positive prostate cancer cells to enzalutamide treatment. As a baseline control, we first evaluated the responses of a panel of AR-positive prostate cancer cells and the AR-negative benign prostate epithelial cells BPH-1 to the TLS inhibitor JH-RE-06. In general, JH-RE-06 at concentrations of up to 2-3 μM was well tolerated, with the overall survival of >50% in comparison with the DMSO control groups (**Fig. 3A**). We then evaluated whether enzalutamide-induced replication stress could render prostate cancer cells particularly vulnerable to TLS inhibition. We found that the presence of JH-RE-06 profoundly sensitized a variety of AR-positive prostate cancer cells, including 22Rv1, TRAMP-C2, LN95, and LNCaP-FGC, to enzalutamide-induced replication stress (**Fig. 3B-E**). In contrast, enzalutamide and JH-RE-06 did not synergize to inhibit proliferation and viability of the AR-negative benign prostate epithelial cell line, BPH-1 (**Fig. 3F**). Therefore, the synergistic effect of JH-RE-06 and enzalutamide on cell viability was strictly dependent on AR expression. Considering the overall lack of response of the AR-negative BPH-1 benign prostate epithelial cells at the highest tested concentrations of enzalutamide (50 μM), JH-RE-06 (3 μM), and their combination, the requirement of enzalutamide-inhibited AR to expose replication stress vulnerability has provided a unique opportunity that could be exploited for targeted treatment of hormone-sensitive prostate cancer though TLS inhibition. Much to our surprise, the synergistic effect of enzalutamide and JH-RE-06 cannot be entirely attributed to the induction of the “BRCAness” state through suppression of HRR genes by AR inhibition (*12*), as TRAMP-C2 cells were already deficient in *BRCA2* (**Fig. S3**), and yet these cells showed a profound synergistic response to the enzalutamide/JH-RE-06 combination treatment (**Fig. 3C**).

**Figure 3.**
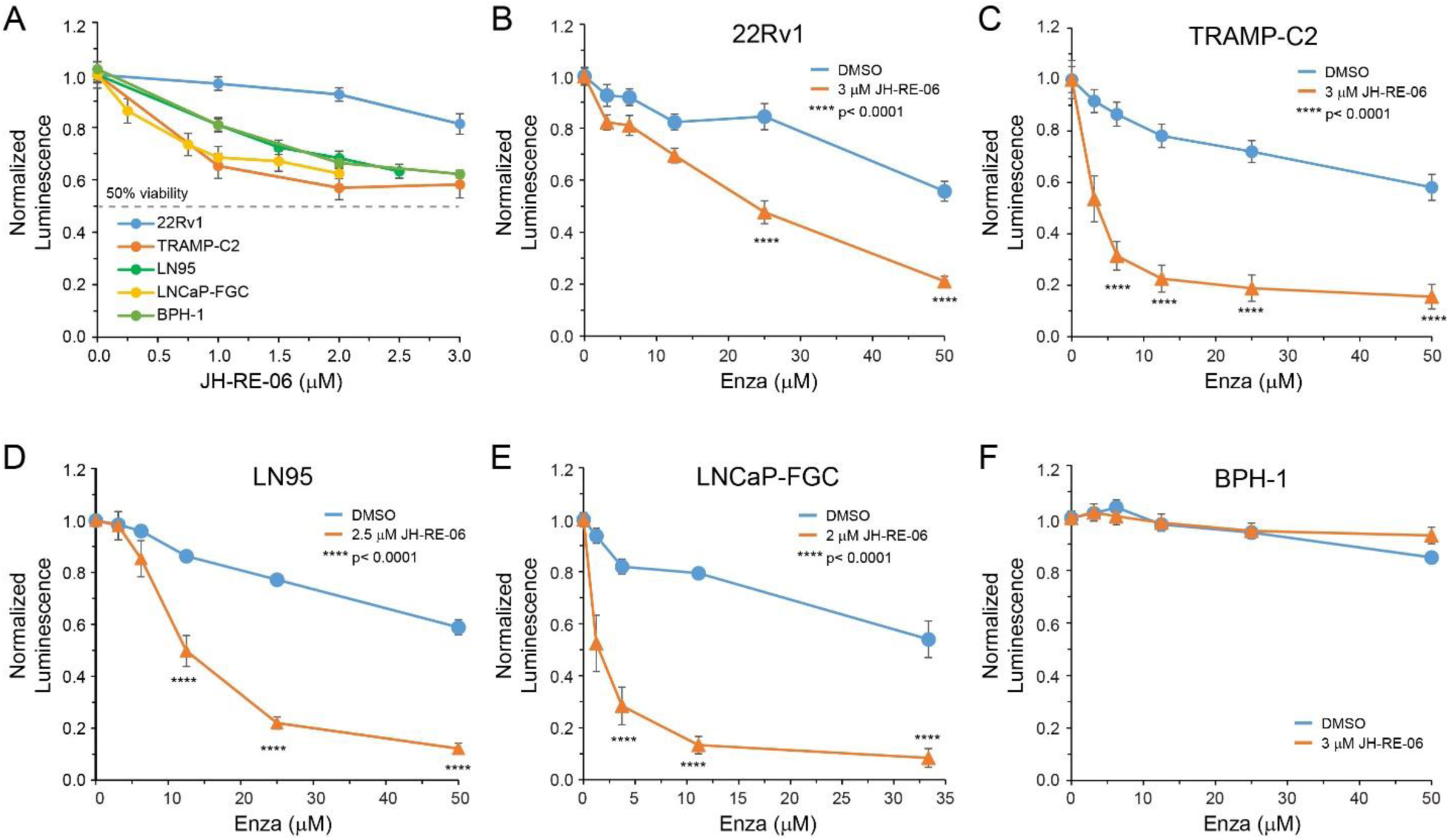
Inhibition of TLS sensitizes AR-proficient prostate cancer cells to enzalutamide-induced replication stress. (A) Dose response of cancerous and benign prostate cells to the TLS inhibitor, JH-RE-06. TLS inhibition by JH-RE-06 sensitizes AR-positive prostate cancer cells, including 22Rv1 (B), TRAMP-C2 (C), LN95 (D) and LNCaP-FGC (E), but not the AR-negative, benign prostate epithelial cells BPH-1 (F), to enzalutamide treatment. Error bars represent S.E.M. Statistical analysis in panels (B-E): student T-test (n=9); ****p<0.0001.

### Systemic Treatment with JH-RE-06 Improves Prostate Cancer Cell Response to Enzalutamide *In Vivo*

Encouraged by the ability of the TLS inhibitor, JH-RE-06, to broadly sensitize a variety of AR-positive prostate cancer lines to enzalutamide treatment, but not in AR-negative benign cells, we employed the TRAMP-C2 murine allograft immune competent tumor model to evaluate whether JH-RE-06 could improve the enzalutamide hormone therapy outcome in a syngeneic *in vivo* model. TRAMP-C2 cells were injected into C57BL/6 mice subcutaneously to grow tumors of approximately 75 mm^3^ size. The mice were randomly distributed into four groups to receive daily intraperitoneal injections of saline, enzalutamide (15 mg/kg), JH-RE-06 (15 mg/kg), and the enzalutamide/JH-RE-06 combination, 5 days a week for 7 weeks. In comparison of the raw tumor volumes (**Fig. 4AB**) and relative tumor volumes normalized to day 0 (**Fig. S4**), we found that individual treatments with enzalutamide or JH-RE-06 had a modest effect to suppress tumor growth, though the difference did not reach statistical significance. In contrast, the combination treatment with enzalutamide and JH-RE-06 resulted in nearly complete inhibition of tumor growth in comparison with the vehicle, JH-RE-06, or enzalutamide treatment groups, in terms of both raw and relative tumor volumes (**Fig. 4 and Fig. S4**), suggesting that co-administration of the TLS inhibitor JH-RE-06 significantly improved the treatment outcome of the enzalutamide hormonal therapy.

**Figure 4.**
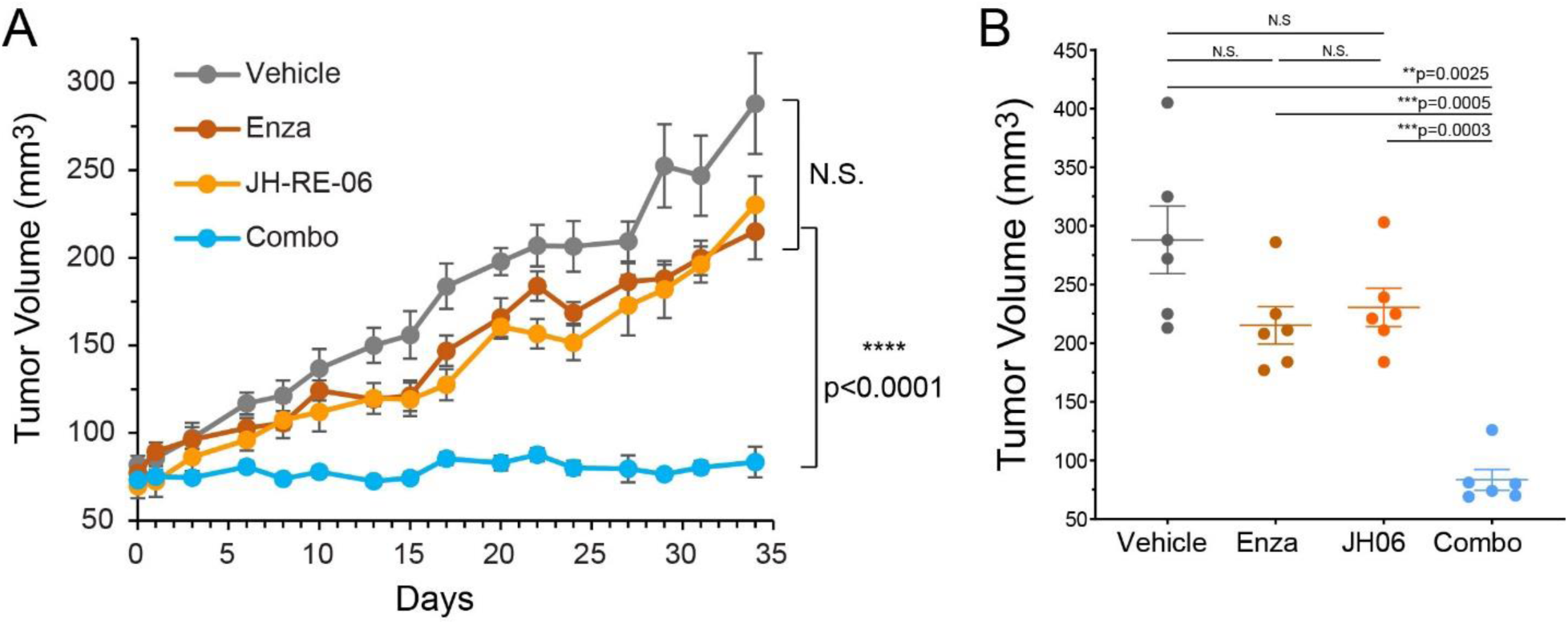
JH-RE-06 improves prostate cancer response to the AR-inhibitor enzalutamide in a syngeneic allograft murine tumor model. (A) Inhibition of TRAMP-C2 allograft tumor growth with intraperitoneal administration of vehicle, enzalutamide, JH-RE-06, and the enzalutamide/JH-RE-06 combination. JH-RE-06 was formulated in 4% methyl-β-cyclodextrin in saline. Enzalutamide was formulated in 24% Captisol, 36% Saline, 32% PEG400, 8% DMSO. (**B**) Tumor values of individual groups at the end of the treatment period. Error bars represent S.E.M. (n=6). Statistical analysis: one-way ANOVA followed by Brown-Forsythe and Welch multiple comparison test. ****p < 0.0001. N.S., not significant.

## DISCUSSION

Nuclear hormone receptor-mediated signaling plays a significant role in fueling the growth of hormone sensitive cancers, such as prostate cancer. Accordingly, inhibition of nuclear hormone receptors, such as AR inhibition by the frontline therapeutic enzalutamide and other ARPIs, can effectively suppress their growth, and this approach has improved overall survival in men with metastatic prostate cancer. However, treatment-induced acquisition of new treatment resistance mechanisms eventually undermines the effectiveness of these therapies. Here, we show that AR inhibition by enzalutamide induces replication stress that relies on TLS to mitigate the accumulation of ssDNA replication gaps. The reliance on error-prone TLS polymerases in this process may be a significant contributing factor to the treatment-induced mutagenesis that may ultimately lead to the inevitable acquisition of resistance. Conversely, the addiction to TLS-mediated resolution of replication stress in enzalutamide-treated AR-positive cancer cells renders them vulnerable to the TLS inhibitor, JH-RE-06. Consequently, TLS inhibition profoundly amplifies the replication stress induced by enzalutamide, and broadly sensitizes a variety of hormone-sensitive prostate cancer lines to enzalutamide *in vitro*. Although intra-tumoral injection of JH-RE-06 has been previous utilized in xenograft tumor models to sensitize melanoma (skin cancer) to cisplatin (*18*) and suppress the tumor growth of lung cancer (*20*) and *BRCA1*-deficient breast cancer (*21*), a systemic delivery of JH-RE-06 would have much broader clinical applications. Here, we demonstrate that systemic delivery of JH-RE-06 with enzalutamide results in striking efficacy in suppressing prostate tumor growth in the murine syngeneic model of TRAMP-C2 cells over vehicle or individual treatments with enzalutamide or JH-RE-06, representing a significant step forward towards eventual clinical applications. As the clinical utility of ARPIs such as enzalutamide is increasingly shifting towards the early stage of prostate cancers that are responsive to hormone therapy before they acquire resistance mutations, the ability of TLS inhibition to sensitize hormone-response prostate cancer to enzalutamide treatment and, at the same time, to suppress the error-prone TLS and mutagenesis may render TLS inhibition a particularly attractive adjuvant therapeutic solution to enhance the therapeutic effect of ARPIs including enzalutamide against hormone-sensitive prostate cancer and prevent or delay the acquisition of new resistance mechanisms leading to castration resistance and the lethal phenotype.

## Acknowledgement

This work was supported by grants from the National Cancer Institute grant R01CA279034 to PZ, JH, JAS, and AJA. CSW would like to thank Gaith Droby, Yang Yang, and Lilly Chiou from the Vaziri lab for valuable inputs on experiments.

## Author Contributions

PZ, CV, JAS, JH, LZ, JLB, AJA, and CSW conceived the project. CSW. conducted (1) in vitro cell viability assay under the guidance of PZ and JZ, (2) biomarker and FACS analysis with the assistance from JRA and under the guidance of CV, (3) time-lapse microscopy under the guidance of JLB, and (4) murine tumor studies with the assistance from JP and under the guidance of JAS, AJA and PZ; CSW prepared samples for RNA-Seq analysis. XZ and JX analyzed the RNA-Seq data under the supervision of DW; XL conducted DNA fiber analysis under the guidance of LZ; HJ, WYL and HL synthesized JH-RE-06 under the guidance of JH; PZ and CSW wrote the manuscript with inputs from all the authors.

## Conflicts of interest

PZ and JH are inventors of a patent on JH-RE-06. The remaining authors declare no competing interests.

## Data and materials availability

All data are available in the main text or the supplementary materials.

## MATERIALS AND METHODS

### Compound synthesis

JH-RE-06, synthesized as described previously (*18*), was treated with 0.1 N NaOH (1 equivalent) in CH_3_OH to yield the corresponding salt form, which was used in this study.

### Cell lines

TRAMP-C2 was cultured in DMEM (Gibco, 11965092), Bovine Insulin (0.005mg/ml) (Sigma, I6634), 5% Nu Serum IV (Corning, 355504), 5% FBS (Cytiva, SH30396.03), 10nM DHEA (Santa Cruz, sc-202573), and 1% Pen/Strep (Gibco, 15140122). 22Rv1 was cultured in RPMI 1640 (Gibco, 11835030), 10% FBS (Cytiva, SH30396.03), 1mM NaPy (Gibco, 11360070), 10mM HEPES (Gibco, 15630080), 4.5g/L Glucose (Sigma, G8769), and 1% Pen/Strep (Gibco, 15140122). LN95 was cultured in RPMI 1640 (Gibco, 11835030), 10% FBS Charcoal Stripped (Sigma, F6765), and 1% Pen/Strep (Gibco, 15140122). LNCaP-FGC was cultured in RPMI 1640 (Gibco, 11835030), 10% FBS (Cytiva, SH30396.03), and 1% Pen/Strep (Gibco, 15140122). BPH-1 was cultured in RPMI 1640 (Corning,10-040-CV), 10% FBS (Cytiva, SH30396.03), and 1% Pen/Strep (Gibco, 15140122). All cells were cultured at 37°C with 5% CO2. Cell lines were tested for mycoplasma contamination with the Lonza’s MycoAlert^®^ Mycoplasma Detection Kit and were confirmed negative.

### RNA sequencing (RNA-Seq) and data analysis

TRAMP-C2 cells were plated on 10 cm dishes and given 24 hours to adhere. Cells were incubated with fresh media containing the appropriate concentrations of either DMSO or enzalutamide (10 μM or 25 μM) for 24 hours. Cells were washed twice with 1X DPBS, collected with a cell scraper to be homogenized via the QiaShredder (Qiagen, 79656) in the RLT buffer, and RNA isolation was performed according to the manufacturer’s protocol using the RNeasy Plus Mini Kit (Qiagen, 74106).

The DDR gene expression was analyzed by Novogene Corporation Inc. (Sacramento CA) and the result visualized using R (version 4.1.0) and the ComplexHeatmap package (version 2.8.0) (*22*). Standardized log2-transformed FPKM values were employed as the expression metrics for the genes represented in the heatmap. Samples were clustered into three groups—DMSO, Enza10, and Enza25—using hierarchical clustering with Euclidean distance, by using the column_split option in the Heatmap function of the CompleHeatmap package. The left annotation of the heatmap indicates the DDR pathway (*23*) associated with each gene. Genes were also clustered hierarchically based on the Euclidean distance of their expression levels. A rotation gene set testing method, ROAST (*24*), from the limma package (*25*) was used with default settings to test whether the set of DDR genes were statistically associated with the treatment groups using random rotations of the data. The p-values for the comparisons between the Enza10 and DMSO groups and between the Enza25 and DMSO groups were both 5 × 10⁻⁴, indicating that DDR genes were significantly associated in the treatment groups (Enza10 and Enza25) in comparison with the control group (DMSO).

### Fluorescence-activated cell sorting (FACS): BrdU incorporation assay

Cells were plated on 10 cm dishes at 50% confluence and allowed to adhere for 24 hours. Media was then replaced with proper supplementation of DMSO, enzalutamide, JH-RE-06, or combo (enzalutamide/JH-RE-06) and treated for 24 hours. Cells were then pulse labeled with 10 µM BrdU for 1 hr and harvested with trypsin-EDTA and fixed in 65% DMEM/ 35% ethanol overnight at 4 °C. Fixed cells were resuspended in 1 ml 2 M HCl for 20 min at room temperature. After centrifugation, the cell pellets were resuspended in 1 mL 0.5 M Borax (pH 8.5) to neutralize any residual acid. Following brief centrifugation, the pellets were washed in 1 mL PBS and then resuspended in 50 µL of antibody labeling solution (30 µL PBS containing 0.5% Tween-20/0.5% BSA plus 20 µL FITC-conjugated anti-BrdU anti-body) (Pharmingen #556028). After 30 min at room temperature in the dark, cells were washed in PBS and resuspended in 1 mL PBS containing 10 µg/mL propidium iodide (PI-Thermofisher P3566) and 8 µg/mL RNase A. The labeled cells were analyzed for PI and BrdU using BD Accuri C6 plus flow cytometer or BD FACSCanto instruments, and data processed using the manufacturer’s software and FlowJo LLC.

### Western blots

To prepare cell extracts containing soluble and CSK-insoluble nuclei, monolayers of cultured cells were washed twice with PBS and lysed in cytoskeleton buffer (CSK buffer: 10 mM Pipes, pH 6.8, 100 mM NaCl, 300 mM sucrose, 3 mM MgCl_2_, 1 mM EGTA, 1 mM dithiothreitol, 0.1 mM ATP, 1 mM Na_3_VO_4_, 10 mM NaF and 0.1% Triton X-100) freshly supplemented with cOmplete protease inhibitor cocktail (Roche) and PhosSTOP (Roche). Lysates were centrifuged at 4000 rpm for 4 minutes to remove the CSK-insoluble nuclei. The detergent-insoluble nuclear fractions were washed once with 0.5 mL of the CSK buffer and then resuspended in a minimal volume of CSK, and then sonication was performed followed by nuclease treatment. For whole cell lysate, soluble and CSK-insoluble fractions were combined, sonicated, and treated with nucleases before analysis by SDS-PAGE and western blotting.

The protein concentrations were normalized using a protein assay dye reagent (Bio-Rad #5000006). Samples were boiled for 5 min at 95 °C and loaded onto SDS-PAGE. After transfer, membranes were blocked with 5% milk in TBST buffer (0.1% Tween-20) for 1 h. Primary antibodies were diluted and incubated overnight, rocking at 4 °C. After three washes with TBST buffer, the secondary antibodies and HRP conjugates were incubated with membranes for 1 h. After three washes, membranes were covered with the ECL reagent and imaged.

### DNA fiber assay

A total of 0.2 × 10^6^ 22RV1 cells were seeded in six-well plates. For experiments involving enzalutamide treatment, enzalutamide (50 μM) was simultaneously added during cell seeding. After 22 hours, cells were labeled with 50 μM CldU for 20 minutes, washed three times with equilibrated culture media, and then labeled with 250 μM IdU for 90 minutes. For experiments involving JH-RE-06 treatment, JH-RE-06 (10 μM) was simultaneously added during the CIdU labeling period. The S1 nuclease DNA fiber assay was performed as published previously (*26*). Briefly, after the two analog labeling steps, the cells were washed twice with 1× phosphate-buffered saline (PBS), followed by incubation in the cytoskeleton buffer (CSK-100) [10 mM MOPS (pH 7), 100 mM NaCl, 3 mM MgCl2, 300 mM sucrose, and 0.5% Triton X-100] for 10 minutes at room temperature to remove the cytoplasm and expose the nuclei. The cells were then gently washed twice with PBS, followed by incubation in the S1 nuclease buffer [30 mM sodium acetate (pH 4.6), 10 mM zinc acetate (pH 6), 50 mM NaCl, and 5% glycerol] containing the S1 nuclease (20 U/mL) at 37 °C for 30 minutes. After removal of the S1 nuclease, nuclei were scraped on ice with 1 mL chilled 0.1% BSA in PBS, spun down at 7000 rpm for 6 minutes at 4°C, and the supernatant was removed, leaving approximately 100 μL of 0.1% BSA in PBS for nuclei pellet resuspension. Two drops, each containing 3 μL of nuclei suspension, were placed near the frosted area of glass slides and air-dried for 2 minutes. Subsequently, 8 μL of lysis buffer [200 mM Tris (pH 7), 50 mM EDTA (pH 8), and 0.5% SDS] were added to each sample drop and incubated at room temperature for 10 minutes. The slides were then tilted at a 15-degree angle to allow the drops to slide down completely to the bottom of the glass slides and air-dried for 10 minutes. Next, the slides were immersed in a 3:1 methanol:acetic acid solution for 10 minutes at room temperature for fixation, followed by three washes in PBS for 5 minutes each. The slides were either stored at 4 °C in PBS or processed immediately. The DNA was denatured in 2.5 N HCl in PBS for 90 minutes at room temperature, followed by three washes in PBS for 5 minutes each. Slides were blocked in the fiber-blocking buffer (2% BSA and 0.1% Tween 20 in PBS) for 30 minutes at 37°C and then incubated with 1:50 rat anti-BrdU (Abcam, ab6326, detect CIdU) and 1:250 mouse anti-BrdU (BD, 347580, detects IdU) for 60 minutes at 37°C. The slides were then washed three times in 0.1% Tween-20 in PBS for 5 minutes each, followed by incubation with 1:100 anti-rat immunoglobulin G (IgG) Alexa Fluor 594 and 1:100 anti-mouse IgG Alexa Fluor 488 for 60 minutes at 37°C. Next, the slides were washed three times in 0.1% Tween-20 in PBS for 5 minutes each, air-dried completely, and mounted with coverslips using ProLong Gold Antifade mounting media. Images were acquired with a Zeiss Axio Observer Z1 microscope using a 63x/1.4 Oil Plan Apochromat DICIII objective and analyzed with Fiji software.

### Cell viability assays

Cells were placed in a 96-well plates with 1000 cells per well and given 24 hours to adhere. Cells were incubated with fresh media containing the appropriate concentrations of JH-RE-06 and enzalutamide for 72 hours. Fresh media was then replenished, and cells were allowed to recover for 48-72 hours before having their luminescence values read. The CellTiter-Glo® 2.0 cell viability assay kit (Promega, G9243) was used according to the manufacturer’s instructions with the substrate being diluted in a ratio of 1∶1. 100 µL of this mixture was added to the cells and mixed on an orbital shaker for 2 minutes. The plates were incubated for 10 minutes at room temperature before luminescence values were measured. The assay was performed in triplicate.

### Time-lapse microscopy

TRAMP-C2 GFP H2B cells were plated at 40k cell density in a 24-well plate using the middle 8 wells to prevent evaporation and allowed to adhere for 24 hours before treatment. Experiments were done in duplicate. Cells were treated with DMSO (negative control), enzalutamide (10 μM), JH-RE-06 (2 μM), and the combination of enzalutamide and JH-RE-06 for 12 hours before conducting live-cell imaging using a Keyence BZ-X810 all-in-one fluorescence microscope equipped with an incubator chamber. Fluorescence and phase-contrast z-stacks were captured at 3-minute intervals for 24 hours using a 20X objective. Best focus projections of the captured images were combined into AVI files. The fates of individual cells were analyzed manually by advancing the AVI files by single frames using ImageJ.

### Syngeneic murine xenograft tumor model

Six-to-eight-week-old, male C57BL/6 mice were purchased from Charles River Laboratories for experimentation. All mice were drug and test naive and were not involved in any previous procedure. All mice were housed in cages in the animal research facility of the Duke University’s Animal Care and Use program, which is fully accredited by the Association for the Assessment and Accreditation of Laboratory Animal Care, International (AAALAC) and meets NIH standards as set forth in the ‘‘Guide for Care and Use of Laboratory Animals’’. The Duke animal facility is maintained under specific pathogen free (SPF) conditions and Duke’s Division of Laboratory Resources performs daily cage checks and maintains feeding and cleaning schedules. The Institutional Animal Care & Use Committee (IACUC) reviewed all animal protocols.

For tumor growth, five million TRAMP-C2 cells were subcutaneously injected into the right flank of six- to eight-week old mice. Tumor growth was monitored by daily palpations to the right flank, and upon adequate tumor size, calipers were used to measure the starting volume. 15 mg/kg enzalutamide formulated in 24% Captisol, 36% Saline, 32% PEG400, 8% DMSO was delivered daily via intraperitoneal injections in the morning, and 15 mg/kg JH-RE-06 formulated in 4% methyl-β-cyclodextrin in saline was delivered daily via intraperitoneal injections in the afternoon. Thrice weekly tumor volume measurements were recorded by calipers using tumor volume = (L/2)*(W^2).

