## Supplementary Materials for "Inhibition of Androgen Receptor Exposes Replication Stress Vulnerability in Prostate Cancer"

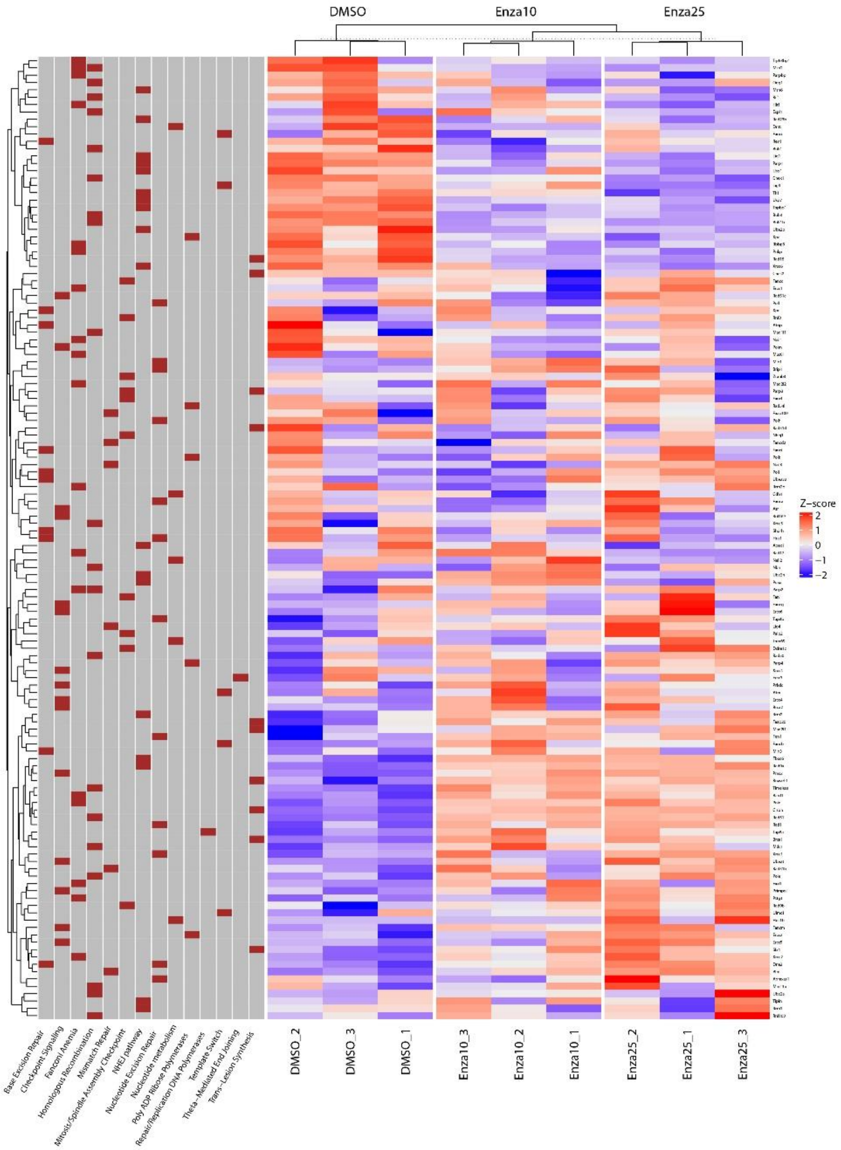


**Supplementary Figure S1.** RNA-Seq analysis revealed a significant perturbation of the transcriptional profile of DNA damage response (DDR) genes in enzalutamide-treated TRAMP-C2 cells in comparison with the DMSO control. Enza10: 10 μM enzalutamide; Enza25: 25 μM enzalutamide. Statistical analysis: gene set testing of DDR genes using the gene set testing method ROAST yields p-values of $5 \times 10^{-4}$ for Enza10 versus DMSO groups and $5 \times 10^{-4}$ for Enza25 versus DMSO groups (threshold: p=0.05).

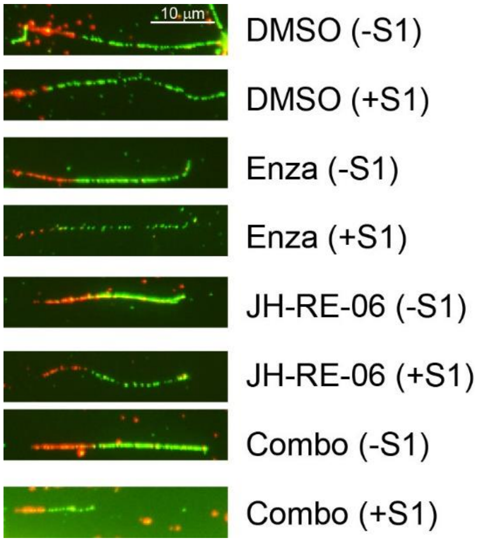


**Supplementary Figure S2.** Representative images of DNA fibers from 22Rv1 cells. Cells were treated with DMSO, 50 µM enzalutamide (Enza), 10 µM JH-RE-06, and their combination, in the absence (−S1) or presence (+S1) of S1 nuclease treatment.

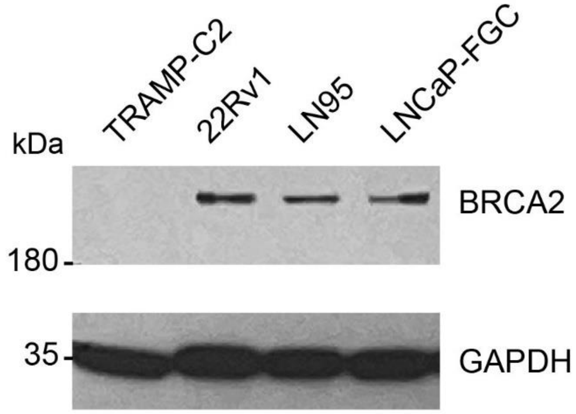


**Supplementary Figure S3.** Western blotting of BRCA2 in selected prostate cancer cell lines.

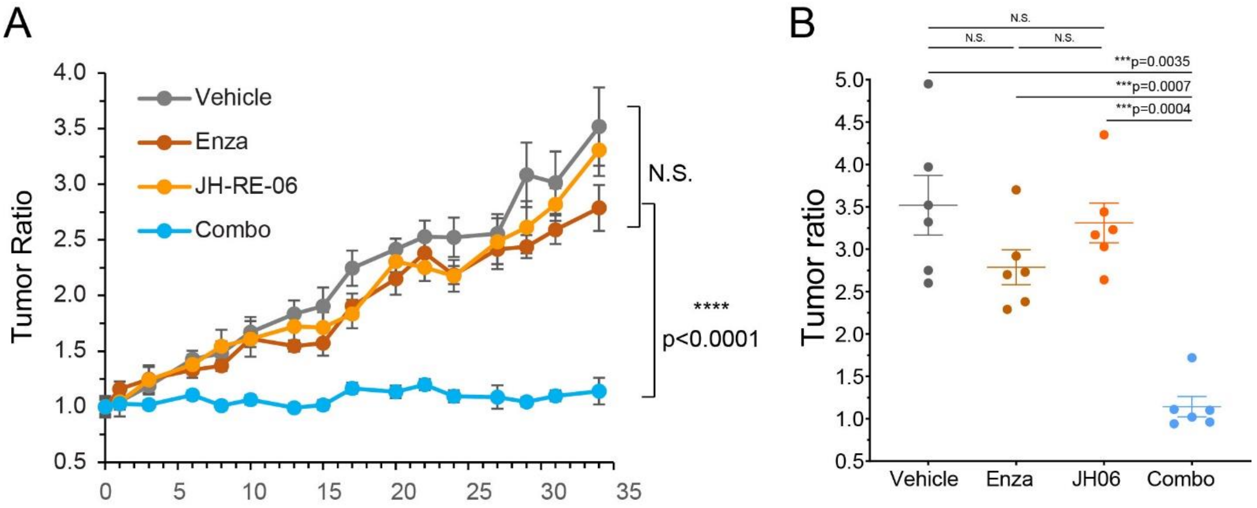


**Supplementary Figure S4.** Relative tumor volume response to the AR-inhibitor enzalutamide and JH-RE-06 in a syngeneic allograft murine tumor model. (A) Inhibition of TRAMP-C2 allograft tumor growth with intraperitoneal administration of saline, enzalutamide, JH-RE-06, and the enzalutamide/JH-RE-06 combination. Tumor ratios were normalized to day 0 of each group to mitigate the potential bias of initial tumor volume variation. (B) Tumor ratios of individual groups at the end of the treatment period. Error bars represent S.E.M. (n=6). Statistical analysis: one-way ANOVA followed by Brown-Forsythe and Welch multiple comparison test. ****p < 0.0001. N.S., not significant.

**Supplementary Table 1. Key Resources**

| Reagent or Resource | Source | Identifier |
|---|---|---|
| Enzalutamide | MedChemExpress | HY-70002 |
| CldU antibody | Abcam | Ab6326 |
| IdU antibody | BD Biosciences | 347580 |
| Goat Anti- Rat Alexa Flour 594 | Invitrogen | A-11007 |
| Goat Anti-Mouse Alexa Fluor 488 | Invitrogen | A-11001 |
| S1 Nuclease (~1200 U/µl) | Thermo Fisher Scientific | 18001016 |
| 5-chloro-2'-deoxyuridine (CldU) – 100MG | Sigma | C6891 |
| 5-iodo-2′-deoxyuridine (IdU) – 5G | Sigma | I7125 |
| Mouse anti- γH2AX S139 clone JBW301 | EMD Millipore | ZMS05636 |
| Rabbit anti-pChk1 S317 | Cell Signaling | 2344 |
| Mouse anti-PNCA (PC10) | Santa Cruz | sc-56 |
| Mouse anti-GAPDH | Santa Cruz | sc-32233 |
| Mouse anti-Vinculin | Sigma Aldrich | V4505 |
| Goat anti-Rabbit | Fortis/Bethyl | A120-101P |
| Goat anti-Mouse | Fortis/Bethyl | A90-116P |
| FITC-conjugated anti-BrdU anti-body | Pharmingen | 556028 |
| Propidium iodide | Thermo Fisher Scientific | P3566 |
| CellTiter-Glo® 2.0 cell viability assay kit | Promega | G9243 |
